# Yap/Taz-TEAD ACTIVITY LINKS MECHANICAL CUES TO CELL PROGENITOR BEHAVIOR DURING HINDBRAIN SEGMENTATION

**DOI:** 10.1101/366351

**Authors:** Adrià Voltes, Covadonga F Hevia, Chaitanya Dingare, Simone Calzolari, Javier Terriente, Caren Norden, Virginie Lecaudey, Cristina Pujades

## Abstract

Cells perceive their microenvironment through chemical and physical cues. However, how mechanical signals are interpreted during embryonic tissue deformation resulting in specific cell behaviors is largely unexplored. The Yap/Taz family of transcriptional co-activators has emerged as an important regulator of tissue growth and regeneration, responding to physical cues from the extracellular matrix, cell shape and actomyosin cytoskeleton. In this work, we unveiled the role of Yap/Taz-TEAD activity as sensor of mechanical signals in the regulation of the progenitor behavior of boundary cells during hindbrain compartmentalization. Monitoring *in vivo* Yap/Taz-activity during hindbrain segmentation we discovered that boundary cells respond to mechanical cues in a cell-autonomous manner through Yap/Taz-TEAD activity. Cell-lineage analysis revealed that Yap/Taz-TEAD boundary cells decrease their proliferative activity when Yap/Taz-TEAD ceased, preceding changes of cell fate: from proliferating progenitors to differentiated neurons. Functional experiments demonstrated the pivotal role of Yap/Taz-TEAD signaling in maintaining the progenitor features in the hindbrain boundary cell population.

## INTRODUCTION

Understanding how cell progenitor specification and differentiation is coordinated alongside morphogenesis to construct the functional brain is one of the challenges in developmental neurobiology. In recent years, there is a growing body of evidences showing that mechanical signals are fundamental regulators of cell behavior. For example extracellular matrix (ECM) rigidity, changes in cell shape and the actomyosin cytoskeleton were found to direct cell behavior in vertebrates through the regulation of the downstream effectors of the Hippo pathway, Yap (Yes-associated protein) and Taz (transcriptional co-activator with PDZ-binding motif, also known as Wwtr1) (Halder *et al*, 2012). A major layer of regulation of Yap and Taz occurs at the level of their subcellular distribution, as the activation of Yap and Taz entails their accumulation into the nucleus, where they bind to and activate the TEAD transcription factors (Zhao *et al*, 2008). Yap and Taz are able to interpret diverse biomechanical signals and transduce them into biological effects in a manner that is specific for the mechanical stress. As example, Yap localization can be regulated by mechanical cues such as ECM rigidity, strain, shear stress, adhesive area or force (Aragona *et al*, 2013; Benham-Pyle *et al*, 2015; Calvo *et al*, 2013; Chaudhuri *et al*, 2016; Dupont *et al*, 2011; Elosegui-Artola *et al*, 2016; 2017; Nakayama *et al*, 2017; Wada *et al*, 2011). Yet, the role of Yap and Taz and their regulation by the multitude of physical tissue deformations occurring during brain morphogenesis remains largely unexplored.

The hindbrain undergoes a dynamic self-organization with dramatic morphogenetic changes during embryonic development, whereby a sequence of mechanical and architectural checkpoints must occur to assess the final functional tissue outcome. This involves the segmentation of the tissue, which leads to the transitory formation of seven metameres named rhombomeres (r1-r7). Rhombomeres constitute developmental units of gene expression and cell lineage compartments (Kiecker & Lumsden, 2005; Fraser *et al*, 1990; Jimenez-Guri *et al*, 2010). This compartmentalization involves the formation of a cellular interface between segments called hindbrain boundary (Guthrie & Lumsden, 1991). Cells at these boundaries express the corresponding rhombomeric markers and exhibit distinct features such as specific gene expression (Letelier *et al*, 2018), and are devoid of proneural genes. In contrast, neighboring rhombomeric regions undergo neurogenesis in a Notch-dependent manner (Nikolaou *et al*, 2009). In addition, boundary cells play very important roles during embryonic development. First, when morphological rhombomeric segments arise boundary cells act as a morphomechanical barrier to prevent cell intermingling. This is due to the enrichment of actomyosin cable-like structures at the apical side of boundary cells that generate tension allowing these cells to behave as an elastic mesh (Calzolari *et al*, 2014). When boundary flanking regions are actively engaged in neurogenesis, hindbrain boundaries constitute a node for signaling pathways instructing the differentiation and organization of neurons in the neighboring rhombomeres (Cheng *et al*, 2004; Riley *et al*, 2004; Cooke *et al*, 2005; Terriente *et al*, 2012). Finally, boundary cells provide proliferating progenitors and differentiating neurons to the hindbrain (Peretz *et al*, 2016). To what extent these distinct functions are intertwined with the cell proliferation and differentiation properties in order to generate a hindbrain with diverse cell types and the correct numbers of cells is still unknown.

We investigated the role of tissue segmentation and mechanical cues in regulating the balance of progenitor vs. differentiated cell state in the embryonic hindbrain, to address how morphogenetic changes were sensed and transduced into specific cell behaviors. We unveiled that Yap/Taz-TEAD activity was restricted to the hindbrain boundary cells acting as a mechanosensor. We revealed by cell-lineage analysis a decrease in the proliferative activity of boundary cells, coinciding with the decline of Yap/Taz-activity within this cell population. This switch in cell proliferation preceded changes in the cell fate, specifically the transition from progenitors to differentiated neurons. Finally, using a combination of functional approaches we demonstrated that Yap/Taz-TEAD activity was essential for maintaining boundary cells as proliferative progenitors. Thus, we propose that mechanical forces within the hindbrain boundaries work as informational systems affecting progenitor maintenance and therefore balancing proliferation and differentiation.

## RESULTS

### Hindbrain boundary cells display Yap/Taz-TEAD activity

During hindbrain segmentation, morphological boundaries are visible as shallow indentations on the outside of the neural tube (Maves *et al*, 2002; Calzolari *et al*, 2014). The hindbrain boundary cells first appeared at the interface between rhombomeres as early as 18hpf (Figure 1A-C; Figure 1-supplementary figure 1A-D’’) marked by *rfng* expression (Terriente *et al*, 2012; Letelier *et al*, 2018), and displayed different morphological features than rhombomeric cells. They differed in their triangular-shape (Gutzman & Sive, 2010) and large apical footprints (Figure 1-supplementary figure 1G, r3/r4 green segmented cells), compared with spindle-shaped rhombomeric cells that have smaller apical sides (Figure 1-supplementary figure 1G, r4 blue segmented cell). In zebrafish, boundary cells actively divided during early embryonic development as observed by BrdU-incorporation (see white arrowheads in Figure 1D), anti-pH3 immunostaining (see white arrowheads in Figure 1E), and live imaging of embryos (Figure 1F, T1-T3 see non-magenta cell incurring into the magenta territory upon mitosis; Calzolari *et al*, 2014). Moreover, boundary cells expressed Sox2 (Figure 1I) supporting the observation that indeed these cells were proliferating progenitors (Galant *et al*, 2016). Interestingly, previous studies showed that boundary cells were not undergoing neurogenesis concomitantly with their rhombomeric cell neighbors; accordingly, proneural genes, responsible of driving neurogenesis, are expressed only in zones that flank rhombomere boundaries (Nikolaou *et al*, 2009).

**Figure 1:**
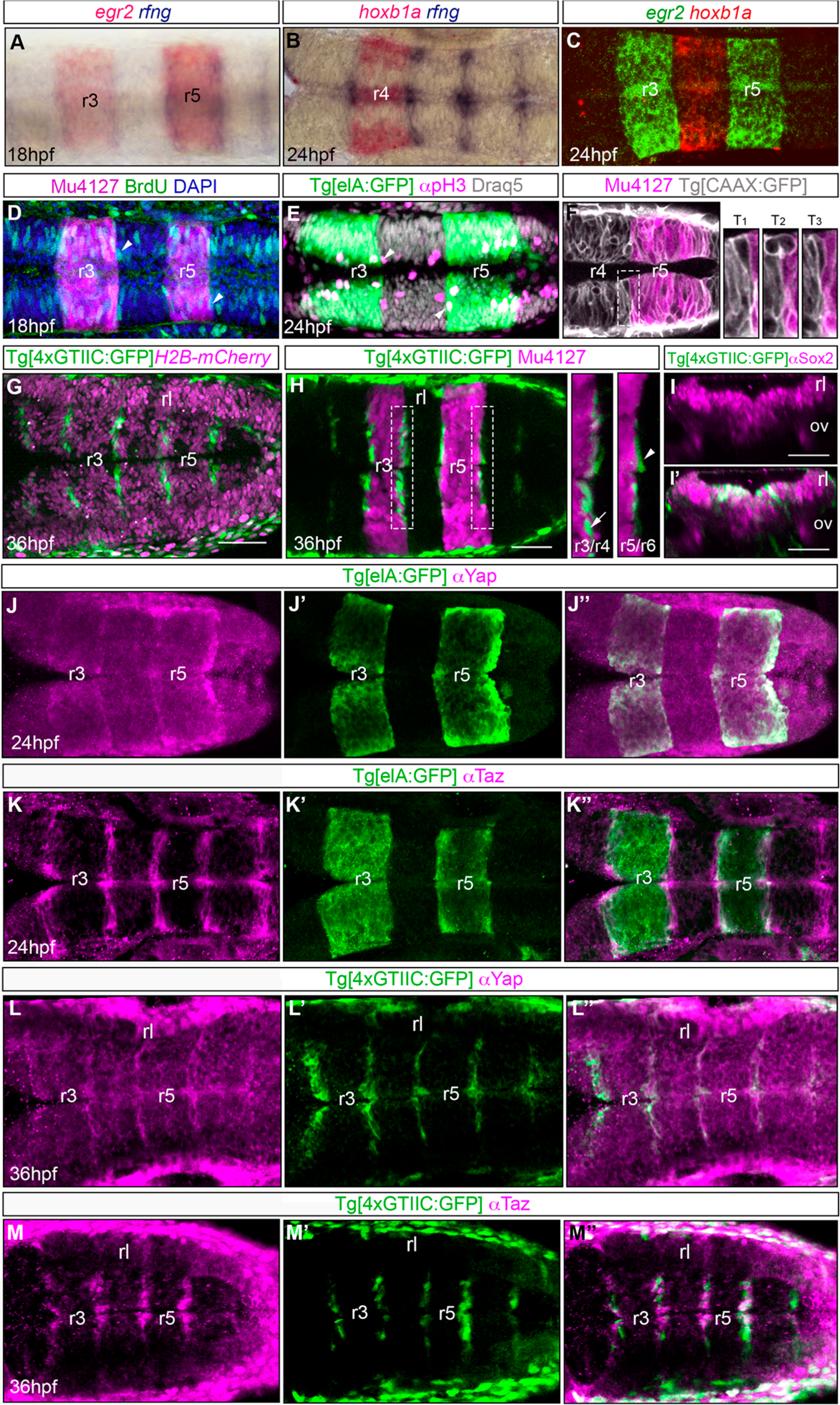
Hindbrain boundary cells display Yap/Taz-activity. A-C) Whole mount double *in situ* hybridization with *egr2a* (A,C), which labels rhombomeres (r) 3 and 5, or *hoxb1a* (B-C) labeling r4, and *rfng* (A-B) labeling hindbrain boundaries in embryos at indicated developmental stages. Note the expression of *rfng* at the interface between adjacent rhombomeres. D) BrdU-staining of a Mu4127 embryo displaying mCherry in r3 and r5 at 18hpf. Note that cells at the boundaries (at the border of mCherry expression shown in magenta) display BrdU-staining in green (see white arrowheads as examples). E) Immunostaining of a Tg[elA:GFP] embryo at 24hpf with anti-pH3 and nuclei stained with Draq5. Note that cells at the boundaries (at the border of GFP expression) display pH3-staining in red/white (see white arrowheads). F) Still from a time-lapse of a double transgenic Tg[CAAX:GFP]Mu4127 embryo displaying GFP in the plasma membrane (shown in white) and mCherry in r3 and r5 (shown in magenta). Inserts from T_1_-T_3_ of the region framed in (F) correspond to different times: note the cell upon division challenges the boundary when it undergoes mitosis. G) Tg[4xGTIIC:GFP] embryo injected with *H2B-mCherry* to visualize nuclei displaying cells with Yap/Taz-activity (green) at 36hpf in discrete progenitor domains of the hindbrain. Note that cells of the rhombic lip (rl) are devoid of Yap/Taz-activity. H) Tg[4xGTIIC:GFP]Mu4127 embryos showing that TEAD-activity is restricted to the boundary cells. Inserts display an enlargement of the r3/r4 and r5/r6 boundaries framed in (H). I-I’) Tg[4xGTIIC:GFP] embryos immunostained with anti-Sox2 (magenta), showing that green Yap/Taz-active cells are located in the ventricular zone and express the cell progenitor marker (I’). J-J’’) Whole mount anti-Yap, and (K-K’’) anti-Taz immunostaining of Tg[elA:GFP] embryo. Note that Yap (J,J’’) and Taz (K-K’’) are preferentially localized in boundary cells (GFP expression border, J’-J’’, K’-K’’). L-L’’) Whole mount anti-Yap, and (M-M’’) anti-Taz immunostaining of Tg[4xGTIIC:GFP] embryos showing that Yap and Taz expression (magenta in L,L’’; M,M’’) overlaps with TEAD-activity (green cells in L’-L’’, M’-M’’). All images are dorsal views with anterior to the left, except for the transverse views in (I’-I’’). r, rhombomere; rl, rhombic lip; ov, otic vesicle.

Boundary cells not only have specific features, but also display distinct functions. We showed that they act as an elastic mesh to prevent cell mixing between adjacent rhombomeres, as a result of the assembly of apical actomyosin cable-like structures that generate tension within this cell population (Calzolari *et al*, 2014). Thus, given the compelling evidence for the relevance of Yap and Taz as downstream mediators of mechanical signals, we wanted to explore whether Yap/Taz-TEAD activity within boundary cells could act as putative sensor of architectural constraints within the hindbrain boundaries and convey this information into a specific cell behavior. First, to determine whether Yap and Taz were acting as activators for TEAD transcription factors in these cells, we monitored TEAD activity *in vivo* by using the transgenic zebrafish reporter line Tg[4xGTIIC:GFP] that expressed GFP under the control of 4 multimerized GTIIC sequences, which are consensus TEAD-binding sites (Miesfeld & Link, 2014). Indeed, Yap/Taz-TEAD activity was confined to discrete territories within the hindbrain (Figure 1G), which coincided with the rhombomeric boundaries since GFP cells were found at the interface between two different rhombomeres (Figure 1H, see inserts with magnifications of r3/r4 and r5/r6). Not only this, boundary Yap/Taz-active cells expressed Sox2 (Figure 1I’), demonstrating that they were neural progenitors. We then performed immunostainings to determine whether the Yap/Taz-TEAD activity was due to Yap or Taz proteins. Although the *yap* transcript is ubiquitously expressed in the embryo (Agarwala *et al*, 2015), the Yap protein was enriched in hindbrain boundaries (Figure 1J-J’’). In addition, Taz protein was restricted to the boundary cells within the hindbrain (Figure 1K-K’’), and as expected, the cells expressing Yap and Taz proteins coincided with boundary cells with TEAD-activity (Figure 1L-M’’). These results suggested that hindbrain boundaries harbored proliferating neural progenitors, enriched in Yap and Taz proteins and displaying TEAD transcriptional activity.

### Establishment of Yap/Taz-TEAD activity in hindbrain boundary cells

To assess the temporal window of Yap/Taz-TEAD activity in the boundary cells, we monitored its onset by following GFP expression in Tg[4xGTIIC:GFP] embryos (Miesfeld & Link, 2014). We found that TEAD-activity started at 20hpf, since transcription of the *gfp* mRNA in the boundaries could be visualized already at this stage (Figure 2A-C), right after hindbrain boundary cells displayed actomyosin-cable structures (15hpf; Calzolari *et al*, 2014). GFP-positive boundary cells were detected slightly later, at 26hpf, and GFP was maintained in the boundaries until 72hpf (Figure 2D-G; data not shown). Next, we unveiled the temporal dynamics of Yap/Taz-activity in the boundary cell population by KAEDE-photoconversion experiments, since KAEDE shifts its emission spectrum from green to red (Ando *et al*, 2002; Hatta *et al*, 2006; Sapède *et al*, 2012). We photoconverted at different times the whole population of KAEDE-Yap/Taz-active cells *in vivo* and traced the photoconverted cell derivatives 18h or 24h later (Figure 2H). To specifically target the Yap/Taz-active cells, we made use of the transgenic line carrying KAEDE under the control of the 4xGTIIC binding-sites (Tg[4xGTIIC:Gal4;UAS:KAEDE]). When green-fluorescent KAEDE (KAEDE^G^) was photoconverted at 30hpf, this resulted in red fluorescent TEAD-active cells born before 30hpf (Figure 2I-I’’’, see no KAEDE^G^-cells in I’). When these very same embryos were observed at 48hpf, the early-born cells were still evident as presence of the converted red-fluorescent KAEDE (KAEDE^R^; Figure 2J, J’’-J’’’), but they also displayed *de novo* synthesized KAEDE^G^ (Figure 2J-J’’’). A thorough analysis to KAEDE^G^-positive cells at 48hpf showed that most of them expressed KAEDE^R^ (compare Figure 2J’ and J’’), indicating that Yap/Taz-TEAD activity in the hindbrain boundaries was triggered before 30hpf as observed in Figure 2A. When KAEDE was completely photoconverted in 48hpf embryos (Figure 2K-K’’’), no KAEDE^G^-cells were present at 72hpf (Figure 2L’) and all the derivatives of Yap/Taz-active cells were KAEDE^R^, suggesting that Yap/Taz-activity was shut off before 48hpf. Thus, our analysis confirmed that Yap/Taz-activity in the hindbrain boundary cells was switched on before 30hpf, just after boundary cells were shown to be important for preventing cell intermingling as a mechanical barrier (Calzolari *et al*, 2014). In addition, these results indicated that the Yap/Taz-activity strongly decreased before 48hpf, even though the GFP was expressed at later stages (Figure 2G), indicating that the Tg[4xGTIIC:GFP] line displayed GFP in boundary cells even beyond the activity declined.

**Figure 2:**
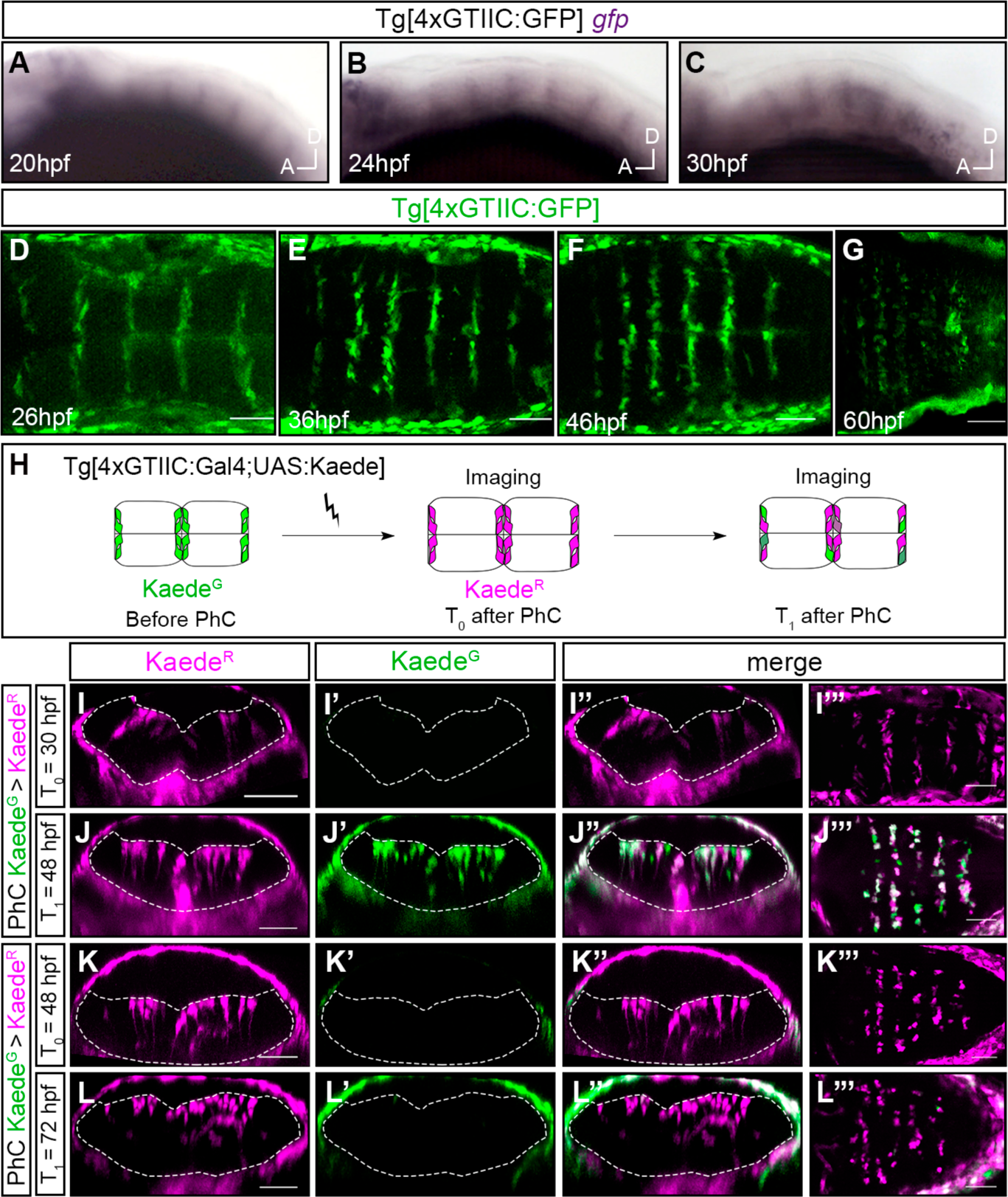
Onset/offset of Yap/Taz-TEAD activity in the hindbrain boundaries. A-C) Tg[4xGTIIC:GFP] embryos assayed for a whole mount *in situ* hybridization with a *gfp* RNA probe at the indicated stages. Note that expression of *gfp* –and therefore Yap/Taz-activity-is already visible in the boundaries at 20hpf. Lateral views with anterior to the left. D-G) Expression of GFP in the hindbrain boundaries in Tg[4xGTIIC:GFP] embryos at indicated stages. Note that GFP is first observed at 26hpf and it is present until 72hpf (data not shown). Dorsal views with anterior to the left. H) Scheme depicting the photoconversion experiment: Kaede^G^ in the hindbrain boundary cells of Tg[4xGTIIC:Gal4FF;UAS:KAEDE] embryos was photoconverted to Kaede^R^ at T_0_, and embryos were let to develop until the desired stage (T_1_). I-J) Embryo in which Kaede^G^ was photoconverted to Kaede^R^ at T_0=_30hpf (I-I’’’) and analyzed at T_1=_48hpf (J-J’’’). Note that new Kaede^G^ is generated in cells between 30hpf and 48hpf (see merged channels in J’’-J’’’). K-L) Embryo in which Kaede^G^ was photoconverted to Kaede^R^ at T_0=_48hpf (K-K’’’) and it was further analyzed at T_1=_72hpf (L-L’’’). Note that no new Kaede^G^-expressing cells are observed after photoconversion (L-L’’’), suggesting the offset of Yap/Taz-activity was before 48hpf. I-I’’; J-J’’; K-K’’; L-L’’) Reconstructed transverse sections of embryos displayed in (I’’’,J’’’,K’’’,L’’’) as dorsal views with anterior to the left.

### Yap/Taz-TEAD activity senses mechanical inputs in hindbrain boundary cells

Changes on the organization of the actomyosin cytoskeleton were found to converge on the regulation of Yap and Taz (for reviews see Halder *et al*, 2012; Panciera *et al*, 2017). Thus, our aim was to address whether integrity of the actomyosin cytoskeleton was necessary for Yap/Taz-activity within the boundary cells. To do so, Yap/Taz-activity was examined after interfering with endogenous tensile forces in the Tg[4xGTIIC:GFP] embryos by several means (Figure 3): i) pharmacological inhibition of myosin II and Rock with *para*-Nitroblebbistatin or Rockout, respectively (Calzolari *et al*, 2014), ii) disruption of the actomyosin cable assembly by downregulating the function of Rac3b -a crucial small Rho GTPase in hindbrain boundaries-with splice-blocking morpholino (Letelier *et al*, 2018), and iii) conditional inhibition of Rac3b by clonal expression of a dominant negative form of Rac3b (Myc:hsp:Rac3bDN; Letelier *et al*, 2018). Upon inhibition of myosin II before the onset of Yap/Taz-activity within the boundaries, the activity was lost as shown by *gfp in situ* hybridization and GFP expression analyses (Figure 3B-C, Blebbistatin: n = 15/20 Rockout: n = 15/19; Figure 3J-K, Blebbistatin: n = 29/42; Rockout: n = 16/31 embryos displaying loss of *gfp*/GFP expression in the boundaries), when compared with control embryos incubated in DMSO (Figure 3A, n = 2/20; and 3I, n = 0/36 embryos with no *gfp*/GFP boundary expression). Both, use of Blebbistatin, which inhibits myosin II by blocking the myosin heads in a complex with low actin affinity (Képiró *et al*, 2014), and use of Rockout that blocks Rho kinase activity (Ernst *et al*, 2012) led to similar results. Similar effects were obtained by downregulating Rac3b with MO-Rac3b (Figure 3H, n = 31/38 embryos lost *gfp* expression in boundary cells), whereas embryos injected with a random morpholino did not display this phenotype (Figure 3G, n = 4/22). In all cases, this downregulation was specific to the boundary cell population, since Yap/Taz-activity within the somites, for example, was maintained (Figure 3B-C, H). Pharmacological treatments neither interfered with boundary cell identity since expression of *rfng* was not affected (Figure 3D-F), nor delayed hindbrain development (Gutzman & Sive, 2010; Calzolari *et al*, 2014). These results showed that tension was necessary for activating the Yap/Taz-pathway; however, it was dispensable for its maintenance since Yap/Taz-activity was not downregulated after inhibiting tensile forces from 26hpf onwards (Figure 3L-N; DMSO: n = 0/16; Blebbistatin: n = 6/24; Rockout: n = 5/19 embryos with loss of boundary-GFP expression). Complementary results were obtained by conditional downregulation of Rac3b and later clonal analysis (Figure 3O): Tg[4xGTIIC:GFP] embryos were injected either with hsp:Myc or Myc:hsp:Rac3bDN, heat-shocked and the percentage of clones within the boundaries that expressed green (Yap/Taz-activity) and red (Myc or Myc:Rac3bDN) was analyzed 16h later (Figure 3P). The majority of boundary clones expressing only Myc displayed Yap/Taz-activity in the control embryos (Figure 3P, Q-Q’’, n = 20/21 see white cells in Q’’). On the contrary, the majority of boundary clones expressing Myc and Rac3bDN did not express Yap/Taz-activity (Figure 3P, R-R’’’, n = 33/47). Thus, Yap and Taz responded to mechanical actomyosin cues, most probably as mediators of mechanical signals.

**Figure 3:**
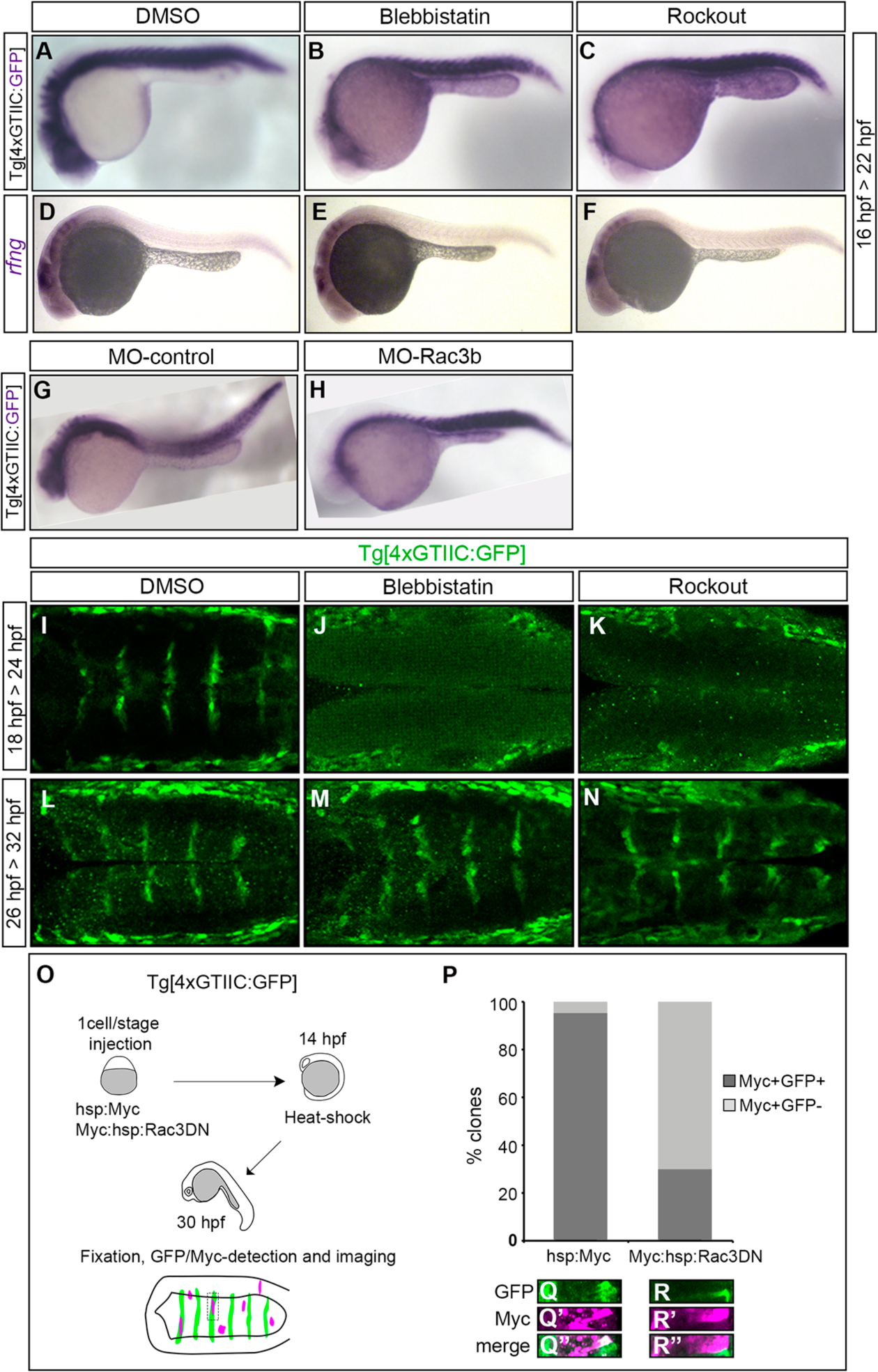
Yap/Taz in the hindbrain boundaries sense mechanical cues. A-C, G-H) Whole mount *gfp in situ* hybridization of Tg[4xGTIIC: GFP] embryos treated with DMSO (A), and myosin II pharmacological inhibitors such as Blebbistatin (B) and Rockout (C) at the indicated interval, or injected with MO-control (G) or MO-Rac3b (H) in order to downregulate Rac3b. Note that in all distinct experimental cases *gfp* expression –and therefore Yap/Taz-activity-is downregulated in the hindbrain boundaries and not affected in the somites. D-F) Whole mount *rfng in situ* hybridization of Tg[4xGTIIC:GFP] embryos treated with DMSO (D), Blebbistatin (E) or Rockout (F). Note that expression of boundary markers is not affected upon treatment as previously shown (Gutzman & Sive, 2010). Lateral views with anterior to the left. I-N) Expression of GFP in Tg[4xGTIIC:GFP] embryos treated with DMSO (I,L), and myosin II inhibitors such as Blebbistatin (J,M) and Rockout (K,N) at different indicated intervals. Note that GFP expression is only disrupted when the treatment is before the onset of Yap/Taz-activity. Dorsal views with anterior to the left. O) Scheme depicting the clonal functional experiment: Tg[4xGTIIC:GFP] embryos were injected with hsp:Myc or Myc:hsp:Rac3DN, heat-shocked at 14hpf and the phenotype was scored at 30hpf. P) Histogram displaying the % of clones with cells expressing GFP (dark grey) of the total Myc-positive cells (light grey), either in control (hsp:Myc, Q-Q’’) and experimental conditions where the actomyosin cables were compromised (Myc:hsp:Rac3DN, R-R’’). Note that when Rac3b is downregulated, the % of Myc-clones with Yap/Taz-activity decreases.

### Yap/Taz-active boundary cells switch the proliferative behavior over time

Next, we studied the spatiotemporal dynamics of Yap/Taz-active cells by exploring their lineage. For this, we established a 4D-imaging pipeline that allowed us to reconstruct cell lineage trees and analyze cell behavior (Movie 1, Figure 4A). We took advantage of the high temporal coverage and resolution provided by Single Plane Illumination Microscopy over multiple embryos encompassing the onset and offset of Yap/Taz-activity (Table 1, Figure 4-figure supplementary 1). Tg[4xGTIIC:GFP] embryos were injected either at one-cell stage with hsp:H2B-GFP and heat-shocked at 26hpf, or at 8 cell stage with H2B-mCherry, and imaged as indicated in Figure 4A and Figure 4-figure supplementary 1. The lineage of 63 single GFP-positive cells expressing RFP/mCherry in the nucleus was reconstructed for an average of 20 hours of imaging (see Movie 1 as example of a single cell tracking), and cell behavior was assessed according to: i) cell division (dividing/non-dividing; Figure 4B-E), and ii) cell differentiation (progenitor/differentiated; Figure 4F-J’’) status. For the analysis of cell proliferation, cells were plotted as a lineage tree in which each line indicated a single cell (Figure 4B). Most of the cells that were tracked from 26hpf onwards actively proliferated as individually depicted as single lines branching upon cell division (orange lines in Figure 4B). Movie 1 and Figure 4C-C’ showed a cell undergoing two divisions and giving rise to four daughter cells. However, from 40hpf onwards cells displayed a clear switch in their proliferative behavior after which most of the tracked cells did not divide any further (black lines in Figure 4B). Although at late stages cells did not express *di novo* GFP because no new activity was triggered (Figure 2), the derivatives of these cells could be tracked due to the stability of GFP. The change in cell proliferative behavior could be observed as well when three KAEDE^G^-cells were photoconverted at 48hpf, and the number of KAEDE^R^-cells remained unchanged 24h later (Figure 4D-D’). Interestingly, a similar behavioral switch was observed when cell proliferation was assessed by an easier approach such as the quantification of the mitotic figures (Figure 4E): the number of Yap/Taz-active boundary cells undergoing mitosis dramatically decreased from 26hpf and 50hpf (19.5% ± 2.3 at 26hpf vs. 5.7% ± 1.4 at 40hpf vs. 2.1% ± 0.5 at 50hpf). On the other hand, adjacent non-boundary cells displayed a different proliferative behavior: the percentage of proliferating *neurog1*- and *atoh1a*-positive cells at 26hpf was half than the boundary cells (Figure 4E; *neurog1*-cells: 8.5% ± 2.6 and *atoh1a*-cells: 11.5%± 1.9, compared with 19.5% ± 2.3 of boundary cells), with a milder decrease over time (*atoh1a*-cells: 5.9% ± 0.9 at 40hpf and 3.2% ± 1.6 at 50hpf; Figure 4E). In summary, the reconstruction of the lineage of Yap/Taz-active boundary cells from *in vivo* data demonstrated that these cells displayed not only a different proliferative activity compared to their neighbors at 26hpf, but they had a proliferation switch at 40hpf coinciding with the decrease of Yap/Taz-activity.

**TABLE 1:**
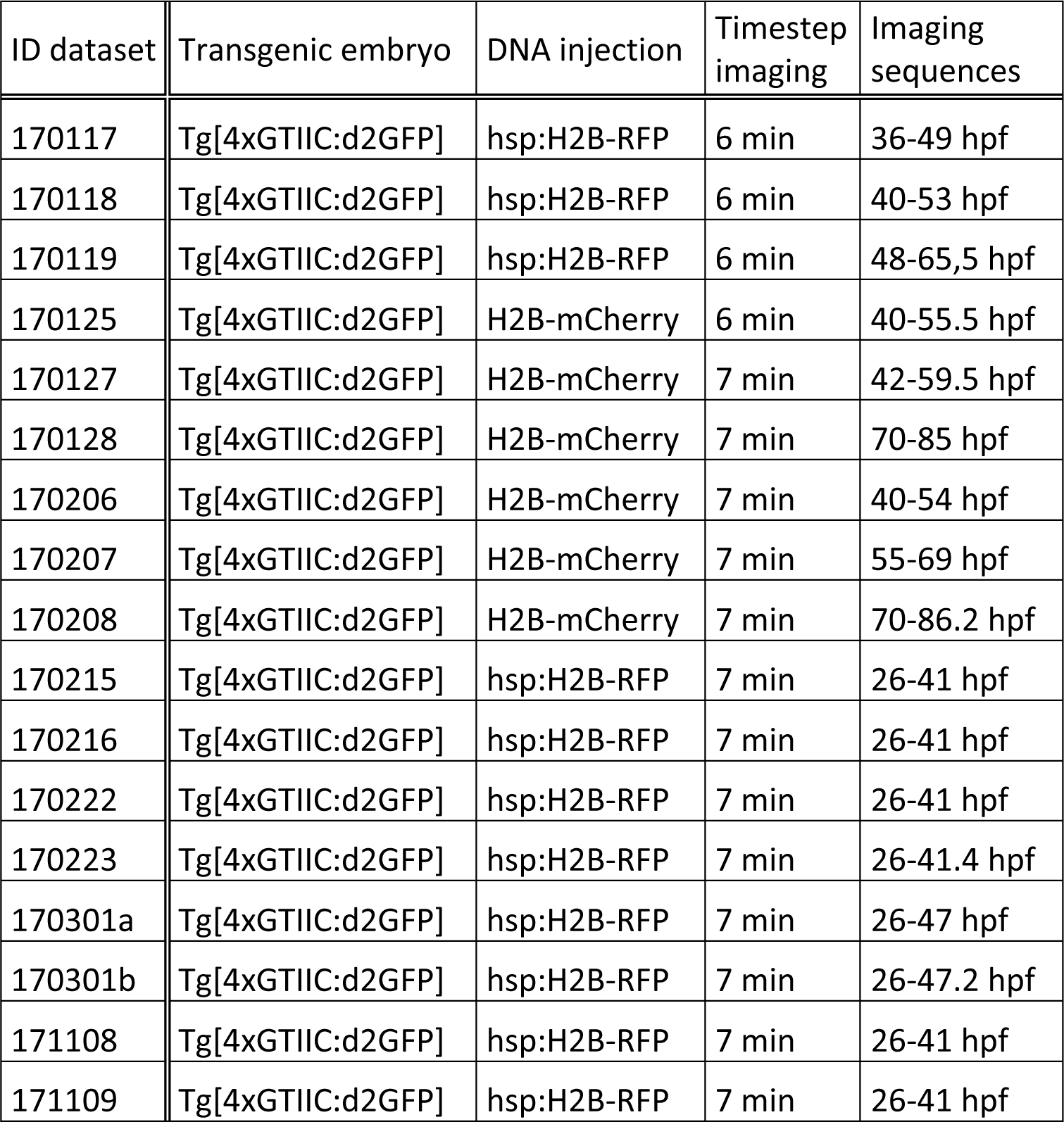
Cohort of embryos and datasets. Datasets used in this study with corresponding information about transgenic embryos and cDNA injections used for Figure 4 experiments. Temporal frequency of image acquisition (timestep imaging) and corresponding imaging sequences are depicted.

**Figure 4:**
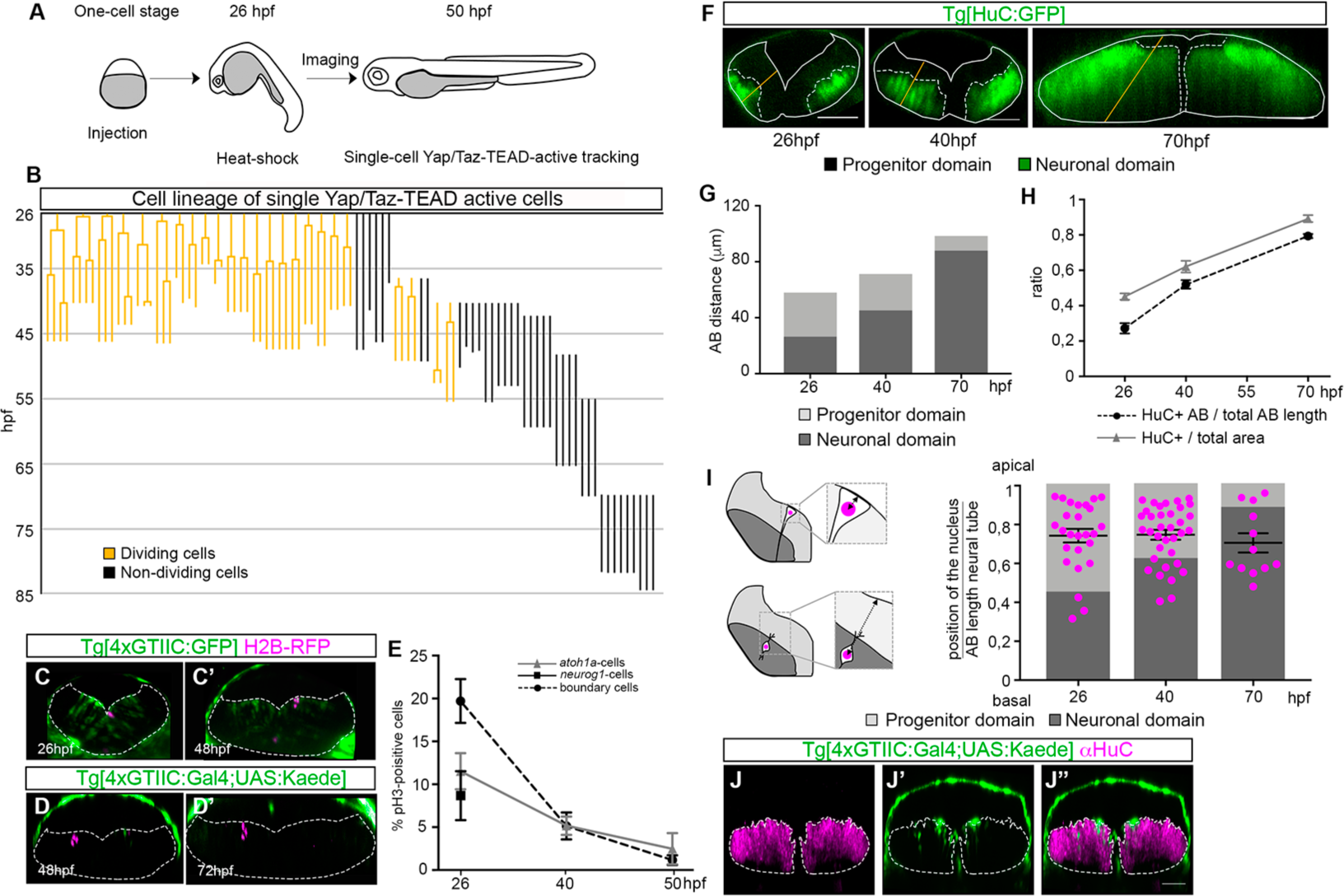
Boundary cells switch their proliferative behavior upon time. A) Scheme depicting the outline of the experiment: Tg[4xGTIIC:GFP] embryos were injected at one-cell stage with hsp:H2B-RPF and heat-shocked at 26hpf, or injected at 8 cell stage with *H2B-mCherry*. Embryos displaying red nuclei within the Yap/Taz-active boundary cells were imaged until the desired stage (Figure 4-figure supplementary 1). B) Representation of the Yap/Taz-active cell lineage tree with Y-axis displaying the time of embryonic development in hours post-fertilization (hpf). The 63 cell lineages are displayed from the moment of tracking onwards and color-coded according to proliferative behavior (orange: dividing; black: non-dividing). Each line corresponds to a single cell starting from the beginning of the movie until the end (Table 1), and branches indicate cell divisions. When the line interrupts it means either the end of the movie or that the cell was lost from the field. Note the switch in cell proliferative behavior from 40hpf onwards, where most of the 4xGTIIC:GFP cells do not branch, thus do not divide. We can track the Yap/Taz-active derivatives beyond Yap/Taz-activity thanks to the stability of the GFP. C-C’) Stills of a time-lapse movie from a single cell followed from 26hpf till 48hpf: the cell undergoes two cell divisions giving rise to 4 cells (Movie 1). D-D’) Example of a group of three cells photoconverted at 48hpf and followed until 72hpf. Note that the number of cells does not change during these 24h, supporting the previous observation of a switch in cell proliferation behavior. E) Graph showing the % of cells undergoing proliferation over time in different hindbrain cell populations: i) Yap/Taz-TEAD proliferating boundary cells (black circle and dashed line); and ii) *atoh1a*-positive rhombomeric cells (grey triangle and solid line) and iii) *neurog1*-positive cells (black square) in the flanking boundary regions. Note the difference in the % of cells undergoing mitosis in boundaries (19%) vs. other hindbrain territories (*atoh1a*-cells: 11.5%; *neurog1*-cells: 8.5%) at 26hpf, and how the boundary cell population dramatically changes the proliferative behavior after the offset of Yap/Taz-activity (5.7% at 40hpf). F) Tg[HuC:GFP] embryos showing the development of the neuronal differentiation domain from 26hpf to 70hpf. G) Histogram displaying the actual size of the progenitor vs. neuronal domains at the indicated stages. Results were obtained from Tg[HuC:GFP] embryos where the apico-basal length was measured (length expanding from the apical ventricular zone edge to the basal mantle zone according to cell orientation within the neural epithelium; n = 8-10 boundaries of 4-5 embryos/stage). Note the dramatic increase in the size of the neuronal domain at expense of the progenitor domain during time. H) Comparison of the ratio obtained from distances between the apical/ventricular and basal/mantle borders of the HuC-domain (black circles and dashed line), with the measurement of the HuC-area (grey triangles and solid line) of an average of 12-18 boundaries (6-9 embryos). Note that both approaches are equivalent for the estimate of the progenitor/neuronal domain progression. I) Analysis of the position along the apico-basal axis of the nuclei of cells tracked in (B) at different times. Positional values were plotted (magenta dots) and overlaid with the information obtained from the progenitor/neuronal map. Briefly, position was scored by measuring the distance of the cell nucleus to the apical side -according to the drawing- and normalized taking into consideration the thickness of the neural tube (see Materials and Methods). Note that most of the cell nuclei at 26hpf lay within the progenitor domain (light grey zone), whereas at 70hpf they are mainly allocated in the neuronal differentiated domain (dark grey zone). J-J’’) Immunostaining of Tg[4xGTIIC:Gal4;UAS:KAEDE] embryos with anti-HuC at 50hpf, showing derivatives of Yap/Taz-active cells within the HuC-positive domain. Although cells within the neuronal domain are not active for Yap/Taz as shown in Figure 2, their derivatives can be tracked due to the stability of the KAEDE protein.

To seek whether this switch in the proliferative behavior of boundary cells was related to the cell differentiation status (progenitor vs. differentiated neuron), we assessed the fate of these very same tracked Yap/Taz-GFP cells by tracing their spatial distribution (position in the ventricular vs. mantle domains) over time. For this, we first generated the dynamic map of differentiated neurons within the hindbrain boundary region (Figure 4F) by following the growth of the differentiation domain (HuC-positive territory) vs. the progenitor domain (HuC-negative territory). A large expansion of the differentiation domain was observed from 26hpf to 70hpf (Figure 4F-G), which required the decrease of the progenitor domain (Figure 4G). The use of the apico-basal (AB) distances for growth assessment was equivalent to the use of HuC-areas, as depicted in Figure 4H. We then plotted the position of the nuclei along the AB axis of the previously tracked cells on the top of the progenitor/differentiation map (Figure 4I). Cell nuclei position was assessed by measuring the distance of each cell nucleus to the ventricular zone at different time steps of the movie (see scheme in Figure 4I). This feature was used as a readout of the cell differentiation state: nuclei located close to the apical side correspond to progenitor cells (light grey, Figure 4I), whereas nuclei close to the basal side correspond to differentiated neurons (dark grey, Figure 4I). Note that at the onset of Yap/Taz-activity, most of the nuclei of tracked cells were found in the progenitor domain (see magenta dots on light grey histogram at 26hpf in Figure 4I). Later on, most of Yap/Taz-derivatives were found within the differentiation domain (see magenta dots on dark grey histogram at 40 and 70hpf in Figure 4I). This switch in nuclei position coincided with the previously observed change in proliferative activity, suggesting that Yap/Taz-active cells within the boundaries behaved as progenitors until they switched off Yap/Taz-activity; then, they ceased proliferating and underwent neuronal differentiation. Most probably boundaries need to balance the ratio progenitors vs. differentiated neurons as other parts of the neural tube do (Hiscock *et al*, 2018). Accordingly, we observed Yap/Taz-derivatives in the neuronal differentiation domain due to the high stability of KAEDE (see green cells within the magenta territory Figure 4J-J’’).

### Yap/Taz-activity regulates cell progenitor behavior in the hindbrain boundaries

To investigate whether Yap and Taz were indeed regulating the proliferative behavior of boundary cells, we knocked-down *yap* or *taz (wwtr1)* and evaluated the effects on cell apoptosis and proliferative activity. First, to dissect the contribution of Yap and Taz, we followed the TEAD-activity in Tg[4xGTIIC:GFP] embryos in which either Yap or Taz were downregulated by splice- or translation-blocking morpholino (Figure 5-figure supplementary 1). No changes on TEAD-activity were obtained upon downregulation of either cofactor suggesting that Yap and Taz function redundantly within the boundary cell population (Figure 5-figure supplementary 1G-I). To further test this possibility, we knocked-down *yap* in *wwtr1* mutants (*wwtr1*^*fu55*^; Figure 5-figure supplementary 1A-D’,J,L). Downregulation of Yap in the *wwtr1*^*fu55/+*^ mutant background had no effect on the number of apoptotic cells, either in the boundaries (Figure 5A; Table 2) or in the rhombomeric cells (Figure 5C; Table 3). Yet, a decrease in the number of proliferating boundary cells was observed (Figure 5B; Table 2), but not within the rhombomeric cells (Figure 5D, Table 3). We made use of *wwtr1* heterozygous mutants (*wwtr1*^*fu55/+*^) injected with MO-Yap, because injection of MO-Yap in *wwtr1*^*fu55*^ homozygous mutant embryos led to early mortality. This observation suggested that Yap/Taz-activity specifically regulated the proliferative activity of hindbrain boundary cells.

**TABLE 2:** Embryos used for functional analysis in the hindbrain boundaries Numbers indicate embryos used for the analysis and the number of boundaries analyzed for Figure 5.

| FIGURE | GENOTYPE | # boundaries | # embryos |
| --- | --- | --- | --- |
| Figure 5A | MO-control *wwtr1*<sup>+/+</sup> | 28 | 7 |
|  | MO-control *wwtr1*<sup>fu55/+</sup> | 108 | 27 |
|  | MO-control *wwtr1*<sup>fu55</sup> | 52 | 13 |
|  | MO-Yap *wwtr1*<sup>+/+</sup> | 36 | 9 |
|  | MO-Yap *wwtr1*<sup>fu55/+</sup> | 88 | 22 |
| Figure 5B | MO-control *wwtr1*<sup>+/+</sup> | 11 | 7 |
|  | MO-control *wwtr1*<sup>fu55/+</sup> | 25 | 13 |
|  | MO-control *wwtr1*<sup>fu55</sup> | 19 | 10 |
|  | MO-Yap *wwtr1*<sup>+/+</sup> | 37 | 19 |
|  | MO-Yap *wwtr1*<sup>fu55/+</sup> | 73 | 37 |
| Figure 5E | *yap*<sup>+/+</sup> *wwtr1*<sup>+/+</sup> | 12 | 3 |
|  | *yap*<sup>fu48/+</sup> *wwtr1*<sup>fu55/+</sup> | 28 | 7 |
|  | *yap*<sup>fu48/+</sup> *wwtr1*<sup>fu55</sup> | 12 | 3 |
|  | *yap*<sup>fu48</sup> *wwtr1*<sup>fu55/+</sup> | 12 | 3 |
| Figure 5F | *yap*<sup>+/+</sup> *wwtr1*<sup>+/+</sup> | 22 | 15 |
|  | *yap*<sup>fu48/+</sup> *wwtr1*<sup>fu55/+</sup> | 45 | 36 |
|  | *yap*<sup>fu48/+</sup> *wwtr1*<sup>fu55</sup> | 23 | 15 |
|  | *yap*<sup>fu48</sup> *wwtr1*<sup>fu55/+</sup> | 22 | 17 |

**TABLE 3:** Embryos used for functional analysis in the rhombomers Numbers indicate embryos used for the analysis and the number of rhombomeres 5 analyzed for Figure 5.

| FIGURE | GENOTYPE | # r5 | # embryos |
| --- | --- | --- | --- |
| Figure 5C | MO-control *wwtr1*<sup>+/+</sup> | 7 | 7 |
|  | MO-control *wwtr1*<sup>fu55/+</sup> | 27 | 27 |
|  | MO-control *wwtr1*<sup>fu55</sup> | 13 | 13 |
|  | MO-Yap *wwtr1*<sup>+/+</sup> | 9 | 9 |
|  | MO-Yap *wwtr1*<sup>fu55/+</sup> | 22 | 22 |
| Figure 5D | MO-control *wwtr1*<sup>+/+</sup> | 6 | 6 |
|  | MO-control *wwtr1*<sup>fu55/+</sup> | 14 | 14 |
|  | MO-control *wwtr1*<sup>fu55</sup> | 10 | 10 |
|  | MO-Yap *wwtr1*<sup>+/+</sup> | 10 | 10 |
|  | MO-Yap *wwtr1*<sup>fu55/+</sup> | 11 | 11 |
| Figure 5G | *yap*<sup>+/+</sup> *wwtr1*<sup>+/+</sup> | 3 | 3 |
|  | *yap*<sup>fu48/+</sup> *wwtr1*<sup>fu55/+</sup> | 7 | 7 |
|  | *yap*<sup>fu48/+</sup> *wwtr1*<sup>fu55</sup> | 3 | 3 |
|  | *yap*<sup>fu48</sup> *wwtr1*<sup>fu55/+</sup> | 3 | 3 |
| Figure 5H | *yap*<sup>+/+</sup> *wwtr1*<sup>+/+</sup> | 13 | 13 |
|  | *yap*<sup>fu48/+</sup> *wwtr1*<sup>fu55/+</sup> | 23 | 23 |
|  | *yap*<sup>fu48/+</sup> *wwtr1*<sup>fu55</sup> | 13 | 13 |
|  | *yap*<sup>fu48</sup> *wwtr1*<sup>fu55/+</sup> | 12 | 12 |

**Figure 5:**
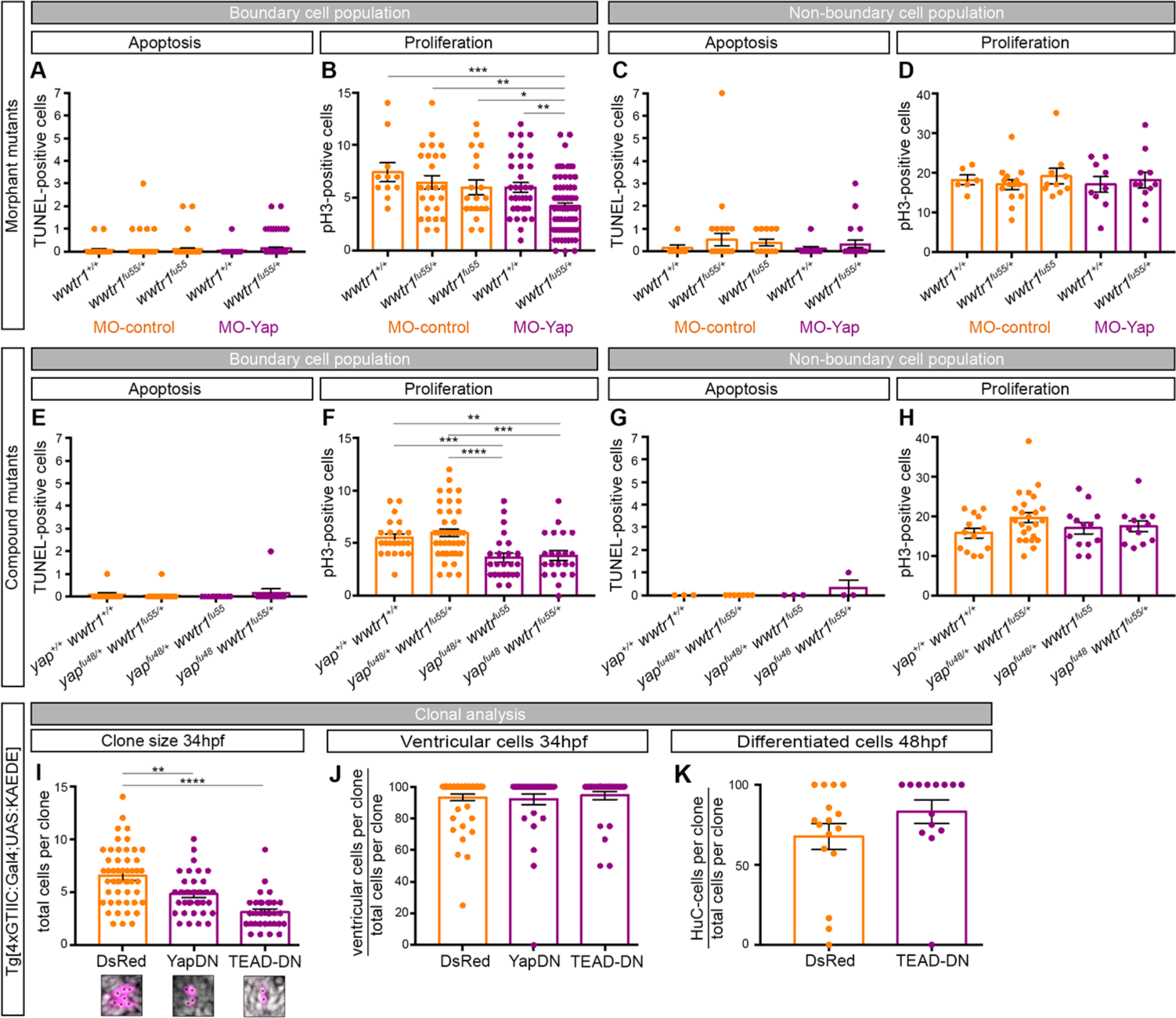
Yap/Taz-activity regulates the proliferative behavior of hindbrain boundary cells. A-D) Wild type (*wwtr1*^*+/+*^), heterozygous (*wwtr1*^*fu55/+*^), or homozygous (*wwtr1*^*fu55*^) embryos for *Taz* mutations were injected with MO-control, and wild type (*wwtr1*^*+/+*^), or heterozygous (*wwtr1*^*fu55/+*^) with MO-Yap, and the number of apoptotic (A,C) and proliferating (B,D) cells within the hindbrain boundaries (A-B; Table 2) or r5 (C-D; Table 3) was quantified at 36hpf. Each dot corresponds to the number of scored cells in a single boundary/rhombomere (see Material and Methods for cell quantification). P values for (B) are the following: MO-control *wwtr1*^*+/+*^ vs. MO-Yap *wwtr1*^*fu55/+*^, p = 0.0008; MO-control *wwtr1*^*fu55/+*^ vs. MO-Yap *wwtr1*^*fu55/+*^, p = 0.0026; MO-control *wwtr1*^*fu55*^ vs. MO-Yap *wwtr1*^*fu55/+*^, p = 0.0361; MO-Yap *wwtr1*^*+/+*^ vs. MO-Yap *wwtr1*^*fu55/+*^, p = 0.0027. E-F) Wild type (*yap*^*+/+*^*wwtr1*^*+/+*^), double heterozygous (*yap*^*fu48/+*^*wwtr1*^*fu55/+*^), or compound mutant (*yap*^*fu48/+*^*wwtr1*^*fu55*^; *yap*^*fu48*^*wwtr1*^*fu55/+*^) embryos for *yap* and *taz* were used for assessing the number of apoptotic (E,G) and proliferating cells (F,H) within the hindbrain boundaries (E-F; Table 2) or r5 (G-H; Table 3). Note that upon dowregulation of Yap in *wwtr1* mutants, and when at least three of the *yap/wwtr1* alleles were mutated, the number of dividing cells decreased within the boundaries. P values for (F) are the following: *yap*^*+/+*^*wwtr1*^*+/+*^ vs. *yap*^*fu48/+*^*wwtr1*^*fu55*^, p = 0.0006; *yap*^*+/+*^*wwtr1*^*+/+*^ vs. *yap*^*fu48*^*wwtr1*^*fu55/+*^, p = 0.0042; *yap*^*fu48/+*^*wwtr1*^*fu55/+*^ vs. *yap*^*fu48/+*^*wwtr1*^*fu55*^, p < 0.0001; *yap*^*fu48/+*^*wwtr1*^*fu55/+*^ vs. *yap*^*fu48/+*^*wwtr1*^*fu55*^, p = 0.0006. Tables 2-3 provide numbers of embryos used for each genotype analysis. Boundary cells were identified as described in Material and Methods. I-J) Tg[4xGTIIC:Gal4;UAS:KAEDE] embryos were injected with UAS:DsRed (DsRed), UAS:DsRed-YapDN (YapDN) and UAS:DsRed-TEAD-DN (TEAD-DN) forms to genetically induce clones expressing DsRed-constructs only in the Yap/Taz-active cells. Embryos with clones were analyzed and the number of cells displaying DsRed in each clone was scored (I), the position of these clones within the ventricular domain was assessed (J) at 34hpf, and the position within the neuronal differentiation domain was analyzed at 48hpf (K). Note that clones in which Yap/Taz-TEAD was downregulated have lower number of cells than the control ones. However, no differences were observed in the differentiation rate. P values for (I) are the following: UAS:DsRed vs. UAS:DsRed-YapDN, p = 0.0021; UAS:DsRed vs. UAS:DsRed-TEAD-DN, p < 0.0001. *p≤0.05, **p≤0.01, ***p≤0.001, ****p≤0.0001. The non-parametric Mann-Whitney test was used.

To confirm this observation, we performed the same analysis by using a combination of *yap/wwtr1* compound mutants (Figure 5-figure supplementary 1J-L). No changes in cell apoptosis within the hindbrain were observed (Figure 5E, G; Table 2), but the number of boundary cells undergoing mitosis was lower in all *yap/wwtr1* compound mutants with three mutated alleles (*yap*^*fu48/+*^*wwtr1*^*fu55*^ and *yap*^*fu48*^*wwtr1*^*fu55/+*^) when compared to the rest of analyzed genotypes (Figure 5F; Table 2). Again, no defects were observed in any of the mutant combinations when non-boundary regions were analyzed (Figure 5H; Table 3).

Finally, to avoid deleterious effects of the mutations and to study whether the observed effect was cell-autonomous, we specifically decreased Yap/Taz-activity within the hindbrain boundary cells by injection of dominant negative forms of Yap and TEAD (UAS:DsRed-YapDN, UAS:DsRed-TEAD-DN; Miesfeld *et al*, 2015) in Tg[4xGTIIC:Gal4;UAS:KAEDE] embryos. Clones expressing DsRed-constructs were evaluated for cell size and position at 34hpf. In all cases in which TEAD-activity was downregulated, either by Yap-DN or TEAD-DN, we observed a decrease in the number of cells per clone (Figure 5I; UAS:DsRed-YapDN = 4.78 ± 0.34 cells/clone; UAS:DsRed-TEAD-DN = 3.12 ± 0.29 cells/clone) when compared to control UAS:DsRed-injected embryos (Figure 5I; 6.54 ± 0.38 cells/clone). This demonstrated that indeed this was a cell-autonomous mechanism. Finally, to address whether the Yap/Taz-mediated regulation of boundary cell proliferation impacted on their transition towards the differentiation state, we followed the position of cell clones along the apico-basal axis upon clonal inactivation of Yap/Taz. No changes in the position of the clones were observed, with most of the cells located in the ventricular zone (Figure 5J; UAS:DsRed-YapDN = 94,52% vs. UAS:DsRed-TEAD-DN = 92,71% vs. control UAS:DsRed = 92,43%). This suggested that Yap/Taz-activity was not involved in transitioning cells to differentiation but mainly controlling the proliferative state of the progenitors. In line with this finding, when the number of differentiated cells was analyzed in UAS:DsRed and UAS:DsRed-TEAD-DN clones at 48hpf, no differences were observed (Figure 5K; UAS:DsRed-TEAD-DN = 72% vs. control UAS:DsRed = 83% of cells in the HuC-domain). These results demonstrated that Yap and Taz are mechanotransducers that regulate the homeostasis of the progenitor pool in the hindbrain boundaries.

## DISCUSSION

In this work, we provide evidence that mechanical inputs lead to Yap/Taz-activity at hindbrain boundaries, which are sites of high mechanical stress during tissue segmentation. Yap and Taz serve as a transducer of physical properties of the microenvironment into a critical cell decision: to remain undifferentiated and maintain the proliferating progenitor pool.

We show that Yap/Taz-activity is confined to the hindbrain boundaries by the use of the Tg[4xGTIIC:GFP] transgenic line as a TEAD-activity reporter. The expression of the transgene not only overlaps with Yap and Taz proteins within the boundaries, but it also matches with the expression pattern of the newly reported zebrafish TEAD reporter line Tg[Hsa.CTGF:nlsmCherry] described as bona fide Yap1/Taz reporter (Astone *et al*, 2018). This suggests that *ctgf* could be one of the targets of this pathway in the boundaries; however, further work is required to reveal the putative targets within the hindbrain. We show that actomyosin-generated tension in boundary cells is most probably the trigger of Yap/Taz-activity to this cell population in zebrafish embryos. Since amniote embryos do not display actomyosin-structures in the hindbrain boundaries (Letelier *et al*, 2018), the mechanism controlling the proliferative capacity of boundary cells might be different, as well as the distinct proliferative capacity between boundaries and rhombomeric regions.

Although chick rhombomeres were described as centers of cell proliferation compared to boundaries (Guthrie *et al*, 1991), other studies proposed that boundaries serve as repetitive pools of stem-like cells at stages when rhombomeres were actively differentiating (Peretz *et al*, 2016). Here, we demonstrate by several means that boundary cells actively proliferate at early developmental stages in zebrafish embryos, exhibiting a distinct proliferative behavior from their rhombomeric neighbors at 26hpf, which were engaged in massive neurogenesis much earlier (Nikolaou *et al*, 2009). Interestingly, when cell proliferation was assayed in HH18 chick embryos (at stages when neuronal differentiation is triggered in boundary cells), the ratio of proliferating cells in rhombomeric vs. boundary regions was 4:1 (Peretz *et al*, 2016). We find that before neuronal differentiation takes place in the hindbrain boundaries in zebrafish (26hpf) the ratio is 1:2, meaning that boundary cells proliferate more than rhombomeric cells. Later on, at the onset of neuronal differentiation within the boundaries (40hpf) the ratio decreases to 1:1, coinciding with the massive growth of the neuronal differentiation domain observed from 40hpf onwards (Figure 4). This suggests that boundary territories change from being an expanding progenitor pool to equate non-boundary proliferative activity and neurogenic capacity. How can we reconcile the different observations in chick and zebrafish? We think that this can be explained by a differential organization of neurogenesis in these two species (Trevarrow *et al*, 1990; Chandrasekhar *et al*, 1997). Zebrafish develops much faster than chick, with a delay of 24-28h between hindbrain segmentation (12hpf) and neuronal differentiation in boundaries (40hpf), when compared with the delay of 40h observed in chick (from HH9.5 to HH18). Thus, since in zebrafish boundary cells undergo neuronal differentiation earlier, the progenitor pool should be expanded in order to maintain the homeostasis and growth of the boundaries, and this expansion is driven by Yap/Taz-activity. Recently, it has been reported that mechanical forces are overarching regulators of Yap/Taz in multicellular contexts for the control of organ growth (Aragona *et al*, 2013).

To shed light on the mechanism that boundary cells use for their functional transitions we have focused on how they switch from the proliferative cell progenitor state to differentiated neurons. We generated compelling insights about the cellular/population dynamics and lineage relationships of the Yap/Taz-active cells and observed that boundary cells dramatically diminish their proliferative activity from 40hpf onwards, coinciding with the downregulation of Yap/Taz-activity. Yap and Taz have been placed as gatekeepers of progenitor activity in other systems. For instance, Yap and Taz are typically found in the nucleus in somatic stem cells or progenitors, being proposed as determinants of stem cell state (Panciera *et al*, 2016), or instrumental for the critical expansion of early progenitor populations in the primitive mesenchyme and overall gut mesenchymal growth (Cotton *et al*, 2017). Accordingly, our functional approaches indicate that indeed Yap/Taz-activity maintains the boundary cells in the progenitor state by controlling their proliferative activity and suggest that levels of mechanical tension and cytoskeletal organization in boundary territories reach the threshold required to activate the transcriptional effects of Yap and Taz. Boundary cells would be maintained in the proliferative progenitor state due to continued Yap/Taz-activity until this activity ceased. Thus, the demonstration that Yap/Taz mechanotransduction can orient cell behavior in hindbrain boundaries highlights the importance of coordinating morphogenesis and cell fate. In line with this, it has been proposed that mechanoactivation of Yap/Taz promotes epidermal stemness in somatic stem cells by regulating Notch (Totaro *et al*, 2017).

In other systems, Yap appears as the main regulator of TEAD-dependent cell functions. For instance, during optic vesicle development, the retinal pigmented epithelium (RPE) fate is compromised in *yap*^*-/-*^ zebrafish embryos, whereas *wwtr1*^*-/-*^ embryos develop normal RPE (Miesfeld *et al*, 2015). However, hindbrain boundary proliferative behavior is not affected either in *yap*^*fu48*^ or *wwtr1*^*fu55*^ embryos, or in Yap- or Taz-morphants, suggesting that both transcriptional co-activators function redundantly. The reasons of the importance of this backup to maintain TEAD-activity are currently unclear. However, one plausible explanation is that they confer robustness to the system because the expansion of the progenitor pool might be a key requirement for the cell population before becoming a proneural cluster domain.

Our results demonstrate that hindbrain boundary cells in zebrafish give rise to two different derivatives: progenitor cells maintained in the ventricular zone and differentiated neurons. It is interesting to speculate that this remaining boundary cell population may behave as long-lasting progenitors, which could be used for development and maturation, or recruited for later events of central nervous system growth or repair. However, to properly address this question further investigation of this new role for boundary cell progenitors is required.

## MATERIALS AND METHODS

### Fish samples

Animals are treated according to the Spanish/European regulations for handling of animals in research. All protocols have been approved by the Institutional Animal Care and Use Ethic Committees and implemented according to European regulations. All experiments were carried out in accordance with the principles of the 3Rs.

Zebrafish (*Dario rerio*) embryos were obtained by mating of adult fish using standard methods. All zebrafish strains were maintained individually as inbred lines. For repairing rhombomeres 3 and 5, two transgenic lines were used: Mü4127, which is an enhancer trap line in which the trap KalTA4-UAS-mCherry cassette was inserted in the 1.5Kb downstream of *egr2a/krx20* gene (Distel *et al*, 2009); and Tg[elA:GFP] that is a stable reporter line where chicken element A from *egr2a* was cloned upstream of the *gfp* reporter (Labalette *et al*, 2011). Tg[neurog1:DsRed] (Drerup & Nechiporuk, 2013) and Tg[atoh1a:Kalta4;UAS:GFP] (Distel *et al*, 2010) label *neurog1*- or *atoh1a*-cells and their derivatives, respectively. Tg[4xGTIIC:d2GFP] line monitors Yap/Taz-TEAD activity (Miesfeld & Link, 2014). Tg[HuC:GFP] line labels early differentiated neurons (Park *et al*, 2000). The mutant lines *yap*^*fu48*^ and *wwtr1*^*fu55*^ were generated using TALEN-induced mutagenesis strategy (Dingare *et al*, 2018).

### Transgenesis

Tg[4xGTIIC:Gal4;UAS:KAEDE] line was generated by injecting one-cell stage Tg[UAS:KAEDE] embryos with the 4xGTIIC:Gal4 vector generated using the Gateway technology (Life Technologies) and the Tol2 kit (Kwan *et al*, 2007). The 4xGTIIC promoter (Miesfeld & Link, 2014) was placed upstream of Gal4FF. One-cell stage embryos were co-injected with a 2nl volume containing 17.5ng/μl of Tol2 transposase mRNA and 15ng/μl of phenol:chloroform purified 4xGTIIC:Gal4 construct. Three or more stable transgenic lines derived from different founders were generated.

### Cell segmentation

For manual segmentation of single cells, ITK-Snap software was used on embryos from Tg[CAAX:GFP]xMu4127 crosses (Figure 1-figure supplementary 1). Single cells located either at the boundary or in the center of the rhombomere were segmented, and the resulting.vtk files were used to display them in FIJI-3D viewer.

### Whole mount *in situ* hybridization

Zebrafish whole-mount *in situ* hybridization was adapted from (Thisse *et al*, 1993). The following riboprobes were generated by *in vitro* transcription from cloned cDNAs: *gfp, hoxb1a* and *egr2a* (Calzolari *et al*, 2014), and *rfng* (Cheng *et al*, 2004). *myl7* and *ephA4* were generated by PCR amplification (*myl7* Fw primer: 5′-GAC CAA CAG CAA AGC AGA CA-3′, *myl7* Rv primer: 5′-TAA TAC GAC TCA CTA TAG GGT AGG GGG CAG CAG TTA CAG-3′; *epha4* Fw primer: 5’-AAG GAG CTA ACT CCA CCG TGC TC-3’ and *epha4* Rv primer: 5’-TAA TAC GAC TCA CTA TAG GGA GAC ATC TGG GTC TTC CTC CAA A-3’) from 24hpf embryos cDNA, adding T7 polymerase binding site at 5′ of the reverse primers and followed by RNA transcription. The chromogenic *in situ* hybridizations were developed with NBT/BCIP (blue) and FastRed (red) substrates. For fluorescent *in situ* hybridization, DIG-labeled riboprobes were developed with fluorescein-tyramide substrate (TSA system). After staining, embryos were either flat-mounted and imaged under Leica DM6000B fluorescence microscope or whole-mounted in agarose and imaged under Leica MZ FLIII stereomicroscope or SP8 Leica confocal microscope.

### BrdU experiments and TUNEL assay

Embryos were incubated with 10μg/μl 5-Bromo-2′-desoxyuridine (Aldrich) for 2 hours prior to fixation. Afterwards they were incubated in 2N HCl for 30 minutes, three times washed in Sodium Borate pH 8.9 and processed for immunohistochemistry. Anti-BrdU BMC9318 antibody (Roche) was used in whole-mount at 1:200.

Distribution of apoptotic cells in the overall hindbrain was determined by TdT-mediated dUTP nick-end labeling of the fragmented DNA. Briefly, after *ephA4 in situ* hybridization embryos were fixed for 30min with 4% (w/v) paraformaldehyde in PBS-Tween. Then, embryos were washed with PBS-Tween before being incubated with TUNEL reaction mixture for 1 hour at 37°C (Roche) followed by PBS-Tween washes. Fluorescein-labeled deoxynucleotides incorporated in apoptotic cells were visualized in a SP8 Leica confocal microscope.

### *In toto* embryo immunostainings

For immunostaining, embryos were blocked in 5%Goat Serum in PBS-Tween20 (PBST) for 1h at room temperature and incubated O/N at 4°C with primary antibody. Primary antibodies were the following: anti-DsRed (1:500, Clontech), anti-GFP (1:200, Torrey Pines), anti-pH3 (1:200, Upstate), anti-Sox2 (1:200 Abcam), anti-Yap (1:100, Santa Cruz), anti-Taz (1:200, Cell Signaling, D24E4), and anti-Myc (1:200, Clontech). After extensive washings with PBST, embryos were incubated with secondary antibodies conjugated with Alexa Fluor^®^488 or Alexa Fluor^®^555 (1:500, Invitrogen). Draq5™ (1:2000, Biostatus, DR50200) or DAPI were used to label nuclei. Embryos were flat-mounted or whole-mounted in agarose and imaged under a Leica SP5 or SP8 confocal microscope.

### Photoconversion experiments

Tg[4xGTIIC:Gal4;UAS:KAEDE] embryos at 30hpf or 48hpf were anesthetized and mounted dorsally in 1%LMP-agarose. KAEDE^G^ was fully photoconverted with UV light (λ=405 nm) using a 20x objective in a Leica SP8 system. Upon exposure of UV light KAEDE protein irreversibly shifts emission from green to red fluorescence (516 to 581). To make sure that all cells within the hindbrain were photoconverted, we did an accurate analysis using confocal microscopy and YZ confocal cross-sections. In the case of the photoconversion of single-KAEDE^G^ cells, embryos at 30hpf expressing mosaic 4xGTIIC:Gal4;UAS:KAEDE were used. In all cases, after photoconversion embryos were returned to embryo medium with phenylthiourea (PTU) in a 28.5°C incubator. At 48hpf or 72hpf, embryos were mounted dorsally and imaged *in vivo* on a Leica SP8 system using PMT detectors and a 20x objective.

### Pharmacological treatments

Treatments with *para*-Nitroblebbistatin and Rockout were applied once the neural tube was already formed to avoid interfering with its early morphogenesis (Calzolari *et al*, 2014). Thus, in all experiments embryos at 14hpf were dechorionated and treated until 20hpf at 28.5°C with: i) myosin inhibitors such as *para*-Nitroblebbistatin (50μM) (Képiró *et al*, 2014) or Rockout (50μM), and ii) DMSO for control experiments. After treatment, embryos were fixed in 4%PFA for further analysis.

### Conditional overexpression

Myc:hsp:Rac3bDN (T71N-mutation) construct was generated by site-directed mutagenesis (QuikChange Site-Directed Mutagenesis Kit, Stratagene #200518), and cloned into the MCS of a Tol2-based custom vector containing a heat shock promoter (hsp) and a Myc-tag (Letelier *et al*, 2018). Tg[4xGTIIC:GFP] embryos were injected at one-cell stage, grown at 28.5°C, and heat-shocked at 14hpf. All embryos were fixed at 18-20hpf, co-immunostained for Myc and GFP, and imaged for further analysis.

UAS:DsRed, UAS:DsRed-YapDN or UAS:DsRed-TEAD-DN constructs were injected in Tg[4xGTIIC:Gal4;UAS:KAEDE] embryos at one-cell stage. Embryos were grown until 24hpf, fixed, Draq5-stained and imaged under the confocal. Size of the clones was analyzed by quantifying nuclei in DsRed-positive boundary clones.

### 3D+time imaging

*Time-lapse movie for in vivo analysis of cell divisions in the rhombomeric boundaries* Anesthetized live double transgenic Tg[CAAX:GFP]Mu4127 embryos were embedded in 1% low-melting point (LMP) agarose with the hindbrain positioned towards the glass-bottom of the Petri dish in order to have a dorsal view with an inverted objective. The video was obtained with an inverted SP5 Leica confocal microscope and processed and analyzed using Fiji software (NIH). Experimental parameters for the video were: voxel dimension (nm): x267.8 y267.8 z629.4, time frame: 10min; total time: 8h; pinhole: 1 Airy; zoom: 2.8; objective: 20x immersion; NA: 0.70.

#### Single-cell tracking experiments

Embryos were anesthetized using 0,04% MS-222 (Sigma) and mounted in 0,6% LMP-agarose in glass capillaries. Time-lapse imaging was performed at 28.5°C on a Zeiss Lightsheet Z.1 microscope. Tg[4xGTIIC:GFP] embryos were injected with hsp::H2B-RFP or *H2B-mCherry* RNA at 1-8cell stage. Embryos injected with hsp::H2B-RFP were heat-shocked for 20min 2h before imaging. The cohort of embryos and datasets used in this study are depicted in Table 1 and Figure 4-figure supplementary 1. Each dataset corresponds to the imaging of a distinct embryo hindbrain. The videos were analyzed, and cells manually tracked using Fiji software (NIH). Experimental parameters were: voxel dimension (nm): x235.5 y235.5 z1000, time frame: see Table 1; total time: see Table 1; zoom: 1; objective: 20x water-dipping; NA: 1.

### Mapping the progenitor and neuronal domains within the hindbrain boundaries

Live Tg[HuC:GFP]xMu4127 embryos were imaged at 26, 40 and 70 hpf under a Leica SP8 confocal. Mu4127 staining was used as a landmark for rhombomeric interfaces. Fiji was used to measure the distance expanding from the apical ventricular zone edge (apical) of the neural tube to the basal mantle zone edge in r3/r4 and r4/r5, and this was called apico-basal (AB) length (black circle and dashed line, Figure 4H). The boundary neuronal domain corresponds to the length encompassing the GFP-expressing territory (dark grey histogram, Figure 4G). On the other hand, the boundary progenitor domain corresponds to the subtraction of the neuronal length to the total distance (light grey histogram, Figure 4G). The temporal dynamics of the ratio (neuronal HuC+ AB length) / (total AB length) was plotted and compared to the ratio (neuronal area) / (whole hemisphere area) (Figure 4H).

The position of the tracked cell nuclei relative to the total AB length was plotted on the top of the progenitor/differentiation map (Figure 4I). Aiming at displaying the data with anatomical coherence, the ratio (position of the nucleus) / (AB length) was subtracted to 1, so values closer to 1 correspond to the cell position in the apical zone, thus the ventricular progenitor domain (Figure 4I).

### Antisense morpholinos

For morpholino knockdowns, embryos were injected at one-cell stage with splicing-blocking morpholino oligomers (MOs) obtained from GeneTools, Inc. MOs were as follows: MO-p53 (Langheinrich et al., 2002), MO-Yap (Agarwala *et al*, 2015), MO-Taz (*wwtr1* gene) 5’-CTG GAG AGG ATT ACC GCT CAT GGT C-3’, MO-Rac3b (see MO-Rac3bSBI4E5 in (Letelier *et al*, 2018). For controls, random 25N morpholino was injected. MO-p53 was included in all MO-injections to diminish putative artifacts (Gerety & Wilkinson, 2011). Efficiency of the injected morpholinos is displayed in Figure 5-figure supplementary 1.

### Phenotypic analyses

#### Morphant mutants and compound mutants

Apoptotic and cell proliferation analyses were performed in 36hpf embryos. Since boundary cells are located at the interface between adjacent rhombomeres and are contributed by two cell-rows, one from each rhombomeres (even/odd-rhombomeres; Figure 1-supplementary figure 1B-D’’), we used as boundary landmarks the borders of *ephA4* expression in r3 and r5. Fluorescent *in situ* hybridization for *epha4* perfectly corresponds with the edges of rhombomeres as shown in Figure 1-supplementary figure 1I-I’’’. Thus, embryos of distinct genotypes were fixed with 4%PFA and assayed for *ephA4 in situ* hybridization prior to cell proliferation or apoptosis analyses. The last row of cells *epha4*-negative and the first row of *epha4*-positive cells (or the other way around depending on the interface) constitute the boundary cell population. Hence, *in situ* hybridization for *epha4* allows the localization of r2/r3, r3/r4, r4/r5 and r5/r6 boundaries. Embryos were imaged under the Leica SP8 confocal. Apoptotic (TUNEL) and proliferating (pH3-expressing) boundary cells were quantified at r3/r4 and r4/r5, whereas r5 was the rhombomeric territory used for non-boundary cell population analysis.

#### Clonal analysis

Injected embryos were fixed with PFA 4% at 34hpf and stained for Draq5. The size of the clones in the boundaries was assessed by quantifying the number of DsRed-positive nuclei. On the other hand, cell fate was analyzed according to cell position in the neural tube, being progenitor cells those in contact to the ventricle. The percentage of ventricular cells per clone was plotted. For cell differentiation analysis, injected embryos were fixed at 48hpf and immunostained for HuC; cells displaying DsRed and HuC were quantified.

## ACKNOWLEDGEMENTS

We are grateful to the referees whose insights improved our paper. We thank Brian Link who kindly provided us with constructs and reagents. We would like to thank S Calatayud and M Linares for technical assistance, and Pujades lab for critical insights. This work was supported by La Marató-TV3 grant 345/C/2014 and Spanish Ministry of Economy and Competitiveness BFU2015-67400-P (MINECO-FEDER, UE) to CP, BFU2016-81887-REDT/AEI, and Unidad de Excelencia María de Maetzu MDM-2014-0370 to DCEXS-UPF. AV is recipient of a predoctoral fellowship from LaCaixa. JT was a recipient of a postdoctoral Beatriu de Pinos fellowship (AGAUR, Generalitat de Catalunya). CP is recipient of an ICREA Academia award (Institució Catalana per la Recerca i Estudis Avançats, Generalitat de Catalunya).

## AUTHOR CONTRIBUTIONS

AV and CP contributed to the concept and design of experiments, and analysis of the results. AV, CFH, CD and CP contributed to the execution of experiments. SC and JT generated tools. CN and VL were involved in the analysis of the results. CP wrote the manuscript.

## SUPPLEMENTARY INFORMATION

**Figure 1 supplementary figure 1:**
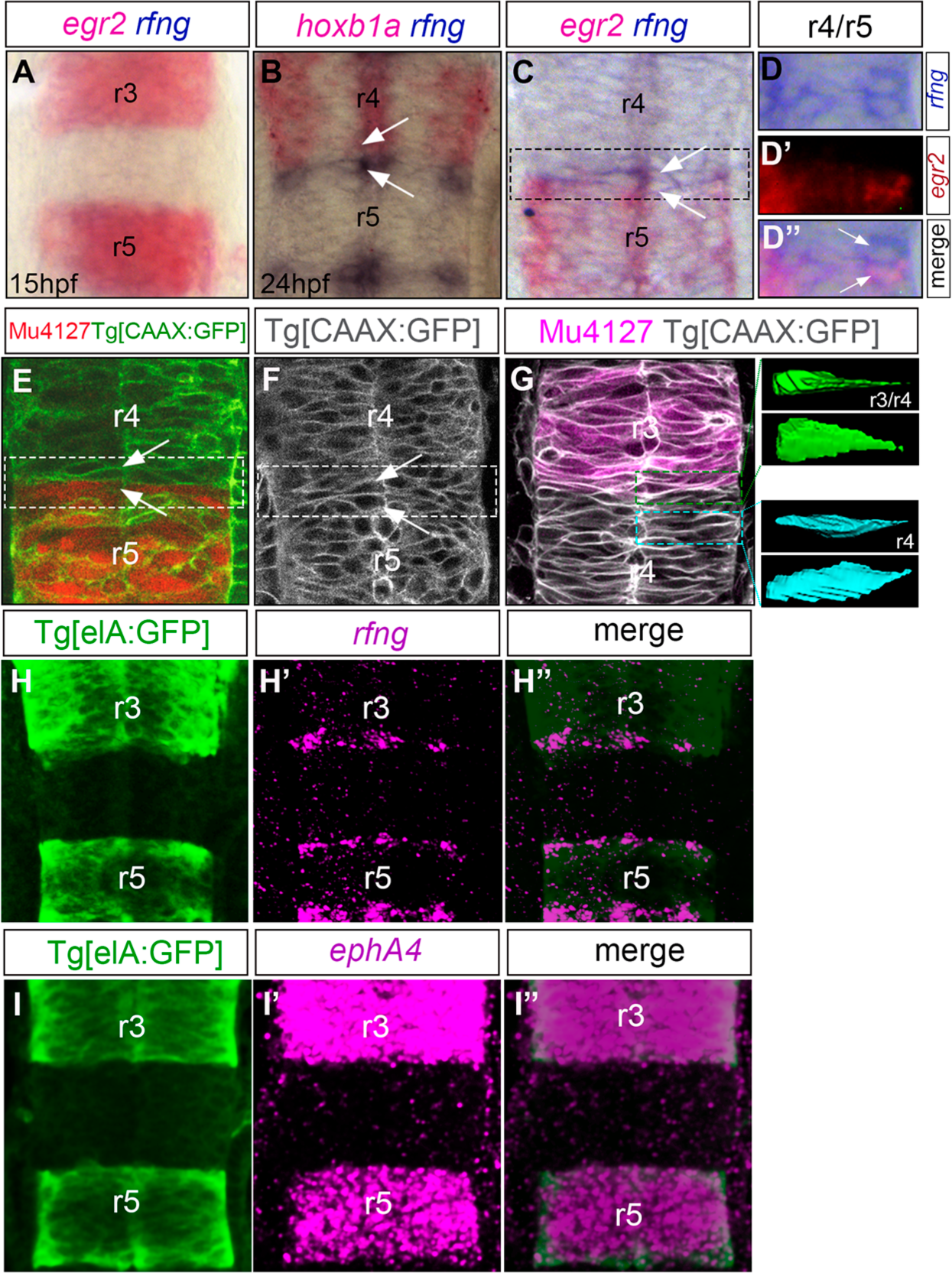
Distinct features of boundary cells. ‘‘) Double *in situ* hybridization for *rfng* -boundary marker-, and either *egr2* -r3 and r5 marker (C-D’’), or *hoxb1a* -r4 marker (B). Note that *rfng* is not yet expressed at 15hpf, and once expressed staining expands two cell rows: one cell row expressing *hoxb1a/egr2a* and *rfng*, and the other one displaying only *rfng* (see cells with white arrows in B-C, and D-D’’). E) Tg[CAAX:GFP]Mu4127 and F) Tg[CAAX:GFP] embryos showing that cells at the boundaries display the singular triangular shape (see white arrows). G) Tg[CAAX:GFP]Mu4127 embryos with inserts from framed regions displaying dorsal (top) and lateral (bottom) views of manually segmented single cells from r3/r4 boundary (green cells) and r4 (blue cells), with the apical side at the left of the image. Note that the green r3/r4 cell has a triangular shape with a large apical side, whereas blue r4 cell is spindle-shaped. H-H’’) Tg[elA:GFP] embryos expressing GFP in r3 and r5 were assayed for *rfng* fluorescence *in situ* hybridization and immunostaining for GFP. Note that *rfng* is expressed at the border of GFP expression, namely the boundary cells. I-I’’) Tg[elA:GFP] embryos expressing GFP in r3 and r5 were assayed for *ephA4* fluorescence *in situ* hybridization and immunostaining for GFP. Note the complete overlapping of *ephA4* and GFP expression, as previously described since *ephA4* is a direct transcriptional target of *egr2*. Dorsal views with anterior to the top. r, rhombomere.

**Figure 4 supplementary figure 1:**
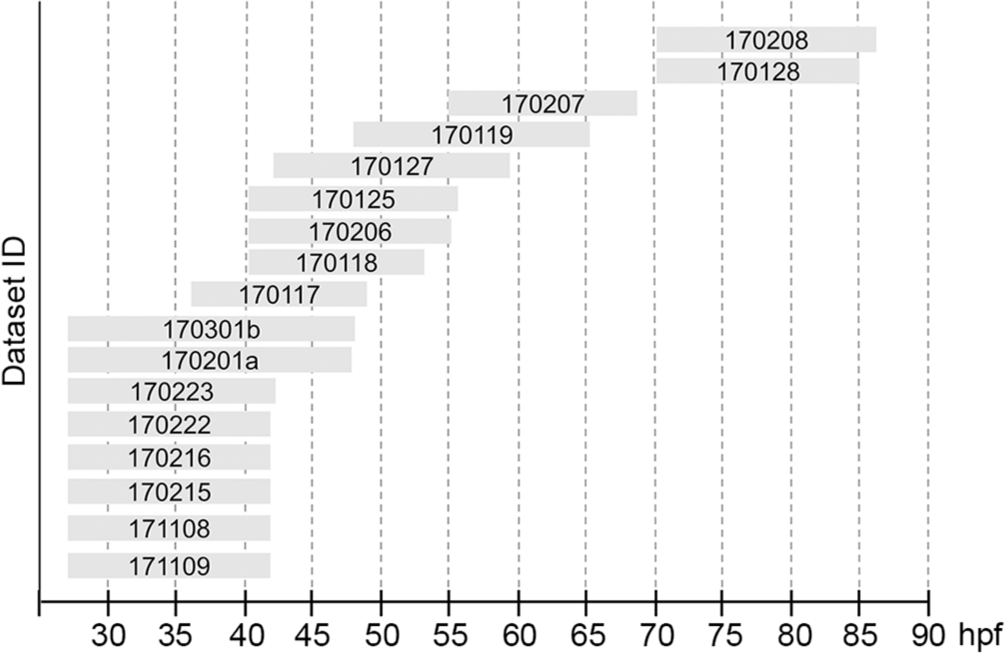
Cohort of embryos and datasets used for the cell lineage studies. Plot displaying the temporal window covering the corresponding dataset.

**Figure 5 supplementary figure 1:**
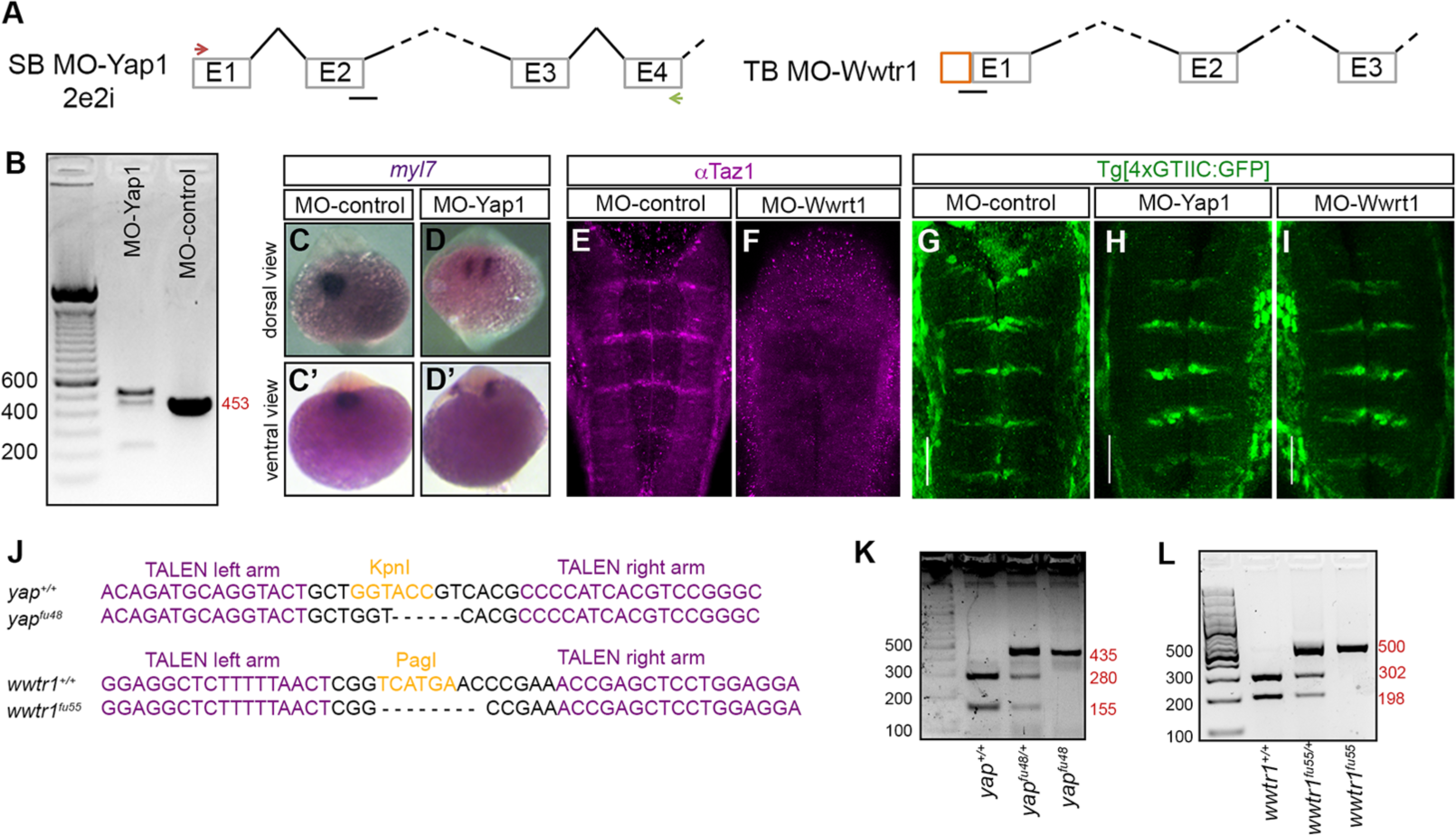
Loss-of-function of Yap and Taz: morpholinos and genomic edition of *yap* and *wwtr1* genes by TALEN technology. A) Scheme depicting the structure of MO-Yap and the position of the splice-blocking morpholino (MO-Yap1SB2e2i) with the corresponding primers to assess its efficiency (see green and red arrows). MO-Wwtr1 is a translation-blocking morpholino for Taz. B) Agarose gel showing the aberrant splicing products induced by MO-Yap as detected by RT-PCR from embryos injected with MO-control and MO-Yap1. C-D) Embryos injected either with MO-control or MO-Yap1 and *in situ* hybridized for *myl7.* Note that upon downregulation of Yap1, embryos displayed defects in the migration of heart progenitors (compare C-C’ with D-D’) as previously described. E-F) Embryos injected either with MO-control or MO-Wwrt1 and immunostained with anti-Taz. Note how Taz protein is downregulated in MO-Wwrt1 injected embryos demonstrating that indeed MO-Wwtr1 interferes with the translation of Taz. G-I) Analysis of Yap/Taz-activity in embryos where Yap or Taz were downregulated using morpholinos (MO-control: n = 16; MO-Yap: n = 23; MO-Wwrt1: n = 26). Note that no effects on Yap/Taz-activity were observed upon downregulation of either cofactor. All images are dorsal views with anterior to the top. J) Alignment of the *yap* (*yap*^*fu48*^) and *wwtr1* (*wwtr1*^*fu55*^) mutant alleles with the corresponding wild-type sequences (*yap*^*+/+*^ and *wwtr1*^*+/+*^) showing: i) the deleted nucleotides, and ii) the TALEN target sites in the *yap* and *wwtr1* loci with the left and right arms (magenta) separated by the spacer including the restriction site used for screening (orange). K-L) Agarose gels showing the obtained bands after enzyme restriction mapping of gDNA corresponding to embryos carrying the wild type or the mutated alleles. Numbers in red indicate the size of the different obtained fragments.

**Movie 1:**
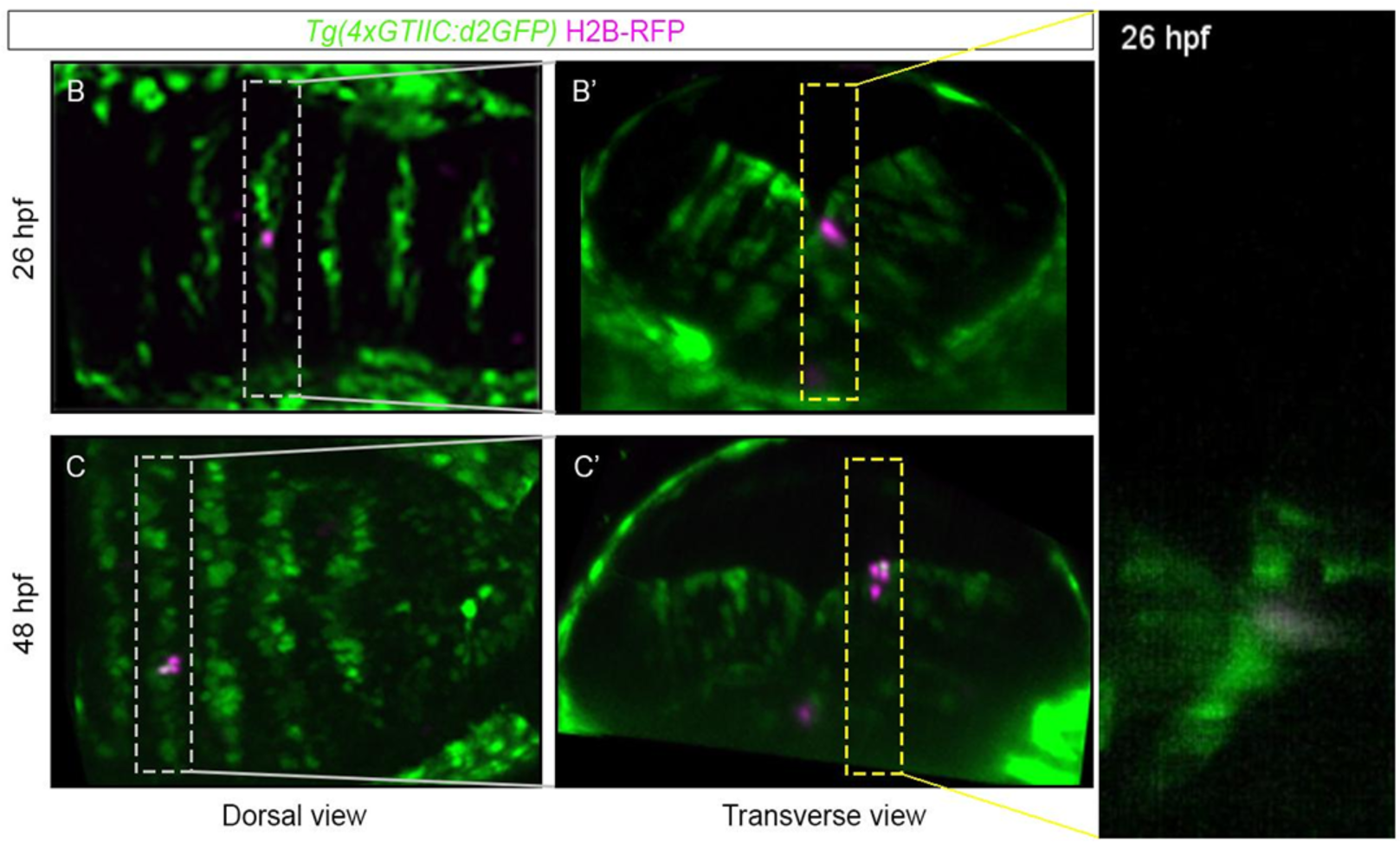
Tracking of a single Yap/Taz-active boundary cell over time. The movie shows how a single red and green cell divides during this time window, and finally gives rise to four daughter cells. All cell lineages depicted in Figure 4B were obtained by tracking single cells as shown in this movie.

